# Restoration of MAIT cell function in healthcare-associated bacterial infections supports recovery of carbapenem efficacy against resistant bacteria

**DOI:** 10.1101/2025.11.21.689870

**Authors:** Xue Yiting, Asad Mustafa Karim, Wan Rong Sia, Fei Han, Kai Lin Chan, Nathalie Grace Chua, Leila Hadadi, Zhenyu Liu, Jeffrey YW Mak, David P Fairlie, Lin-Fa Wang, Johan K Sandberg, Andrea Lay-Hoon Kwa, Edwin Leeansyah

## Abstract

Antimicrobial resistance (AMR) presents a major clinical challenge to patients with healthcare-associated infections (HAIs), particularly among immunocompromised individuals and patients with comoribidites, who often exhibit an impaired mucosa-associated invariant T (MAIT) cell pool. MAIT cells are innate-like T cells enriched in mucosal tissues, possess potent antibacterial activity, and restoration of their function may offer a host-directed strategy against drug-resistant pathogens. We evaluated how stimulation with cognate antigen in combination with various cytokines, modulates MAIT cell cytotoxicity and enhances carbapenem activity. Under optimal conditions, MAIT cells exhibited increased expression of antimicrobial cytolytic proteins and efficiently killed cells pulsed with MAIT cell antigen. IL-15 or IL-2 plus IL-7 were particularly effective in promoting polyfunctional cytotoxic responses. Secretomes from cytokine-stimulated MAIT cells restored the activity of imipenem against engineered *E. coli* expressing the clinically relevant carbapenemases *bla*_NDM-1_, *bla*_KPC-2_, and *bla*_OXA-48_, strongly reducing bacterial growth, viability, and metabolic activity. Notably, IL-2 plus IL-7 stimulation enabled expansion and functional restoration of MAIT cells from HAI patients, whose baseline MAIT cell numbers and responses were diminished. These findings demonstrate that tailored stimulation can reinvigorate MAIT cell effector function and augment antibiotic efficacy, supporting a role for MAIT cells in adjunct immunotherapeutic strategy to combat AMR in vulnerable patient populations.

Category of manuscript: Research Article.

## INTRODUCTION

Antimicrobial resistance (AMR) is a significant global public health crisis, with gram-negative bacteria (GNB) identified as major contributors to healthcare-associated infections (HAIs) [1,2]. The World Health Organization (WHO) has specifically highlighted carbapenem-resistant (CR) Enterobacterales as priority pathogens for urgent research and development of new antimicrobials [3]. Carbapenemases produced by these bacteria degrade one of the last-resort antibiotics, the carbapenems, severely limiting treatment options. Given the plasmid-mediated nature of carbapenemase expression, this resistance can spread rapidly among gram-negative bacteria, particularly in healthcare settings [4]. Therefore, there is an urgent need for innovative and alternative strategies, such as host-directed approaches that harness antibacterial immune cells.

Mucosa-associated invariant T (MAIT) cells are innate-like T cells characterized by a semi-invariant T cell receptor (TCR), primarily composed of the Vα7.2 chain paired with Jα33, Jα20, or Jα12 gene segments [5–7]. This unique TCR configuration enables MAIT cells to recognize vitamin B2 (riboflavin) metabolites derived from bacterial biosynthesis pathways presented by the major histocompatibility complex class I-related protein (MR1) [8,9]. MAIT cells constitute up to 10% of circulating T cells and are enriched in mucosal tissues, including the human gut [10–13]. Because the riboflavin biosynthetic pathway is broadly conserved among bacteria, including in GNB [9,12,14], MAIT cells can detect riboflavin-derived antigens from diverse microbes via TCR–MR1 recognition [8,15–17]. Upon activation, MAIT cells rapidly secrete pro-inflammatory cytokines and antimicrobial cytolytic proteins, including perforin (Prf), granulysin (Gnly), and granzymes (Grz) A and B [15,18,19]. The antimicrobial response of MAIT cells is enhanced by elevated cytokine levels present within the infection microenvironments, including IL-2, IL-7, IL-12, IL-15, IL-18, and IL-23, which increase their sensitivity to low-level TCR signals from bacterial antigens and modulate their activation, proliferation, and effector functions [10,20–25].

MAIT cell–derived cytolytic proteins can eliminate bacteria through several complementary mechanisms. Gnly disrupts bacterial membrane integrity and contributes substantially to MAIT cell–mediated antimicrobial activity [4,18,26]. In conjunction with Gnly, GrzB can kill both intracellular and extracellular bacteria, partly by promoting oxidative damage [18,27,28]. These cytolytic mediators also increase bacterial membrane permeability, which may enhance carbapenem uptake even in strains expressing carbapenemases, while simultaneously exerting direct bactericidal effects independent of antibiotic action [18]. Through these combined mechanisms, MAIT cell cytolytic proteins have the capacity to eliminate carbapenem-resistant bacteria and restore carbapenem efficacy [18].

Individuals and patients with comorbidities are groups disproportionately affected by AMR infections. This key population often exhibits reduced MAIT cell frequencies and impaired antimicrobial responses [4,11,29–33]. This dysfunction may contribute to their increased susceptibility to drug-resistant infections and poorer clinical outcomes. Enhancing MAIT cell responses in these populations could, therefore, represent a valuable immunotherapeutic strategy. Here, we show that combined TCR- and cytokine-mediated activation shapes MAIT cell cytotoxicity, with key cytokines modulating effector functions in distinct, time-dependent manners. Under these defined conditions, MAIT cells acquired an enhanced antimicrobial profile capable of eliminating carbapenem-resistant *Escherichia coli* and restoring carbapenem activity. Notably, this approach also expanded and functionally restored MAIT cells that were numerically and functionally compromised in HAI patients. Collectively, these results outline a therapeutic framework in which cognate antigen and cytokine-based activation can enhance MAIT cell immunity and strengthen host defense against antimicrobial-resistant pathogens, particularly in vulnerable or immunocompromised individuals.

## RESULTS

### Reduced MAIT cell frequency and antimicrobial effector function in patients with healthcare-associated infections

Hospitalized patients are highly susceptible to HAIs, many of which involve multidrug-resistant organisms. Because MAIT cells play an important role in antimicrobial defense, we examined their frequency and effector profiles in a cohort of HAI patients (S1 Table). MAIT cell levels were significantly reduced in HAI patients (Figure 1A) with a concomitant increase of active caspase 3 expression (Figure 1B). This reduction did not correlate with age, suggesting the decline in MAIT cell numbers in the current cohort was age-independent (S1 Figure). The remaining MAIT cells displayed elevated levels of the cytolytic proteins GrzB and Prf (Figure 1C). However, expression levels of GrzA and Gnly remained unchanged (Figure 1C). Interestingly, MAIT cells showed impaired degranulation and cytolytic capacity as well as pro-inflammatory cytokine expression following bacterial stimulation (Figure 1D – 1G). Together, these findings indicate that MAIT cells in HAI patients are both numerically and functionally impaired, showing reduced antimicrobial responsiveness to bacterial stimulation despite an activated resting phenotype.

**Figure 1.**
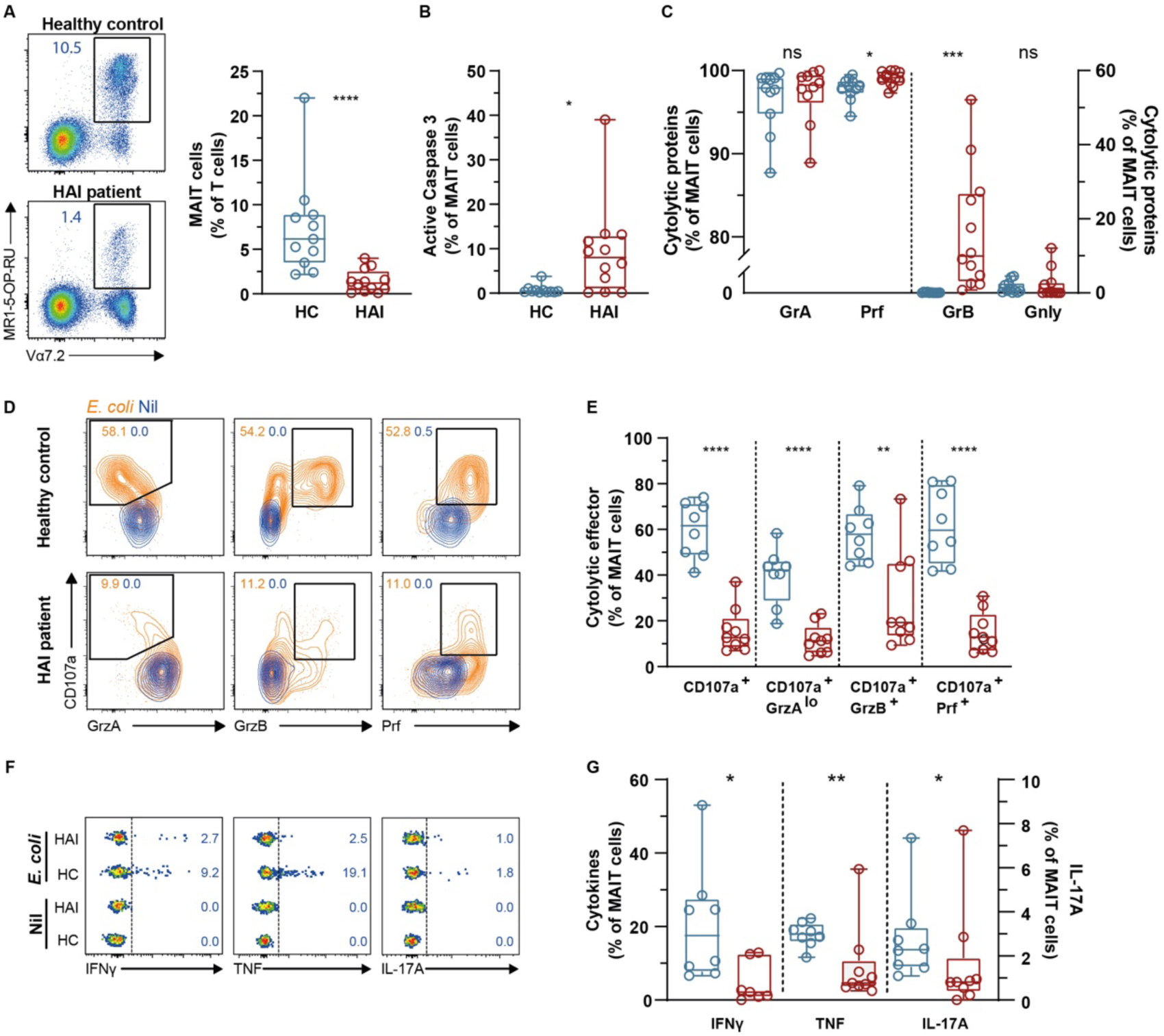
Depletion of MAIT cells in HAI patients with defective effector function following *E. coli* stimulation. **(A)** MAIT cell frequency in peripheral blood of healthy donors (n=11) and HAI patients (n=12) identified as Vα7.2^+^MR1-5-OP-RU^+^CD3^+^ T cells. (**B, C**) Expression of active Caspase 3 **(B)**, GrzA, GrzB, Gnly, and Prf **(C)** within MAIT cell population. **(D, E)** MAIT cell cytolytic effector function in healthy donors (n=5) and HAI patients (n=7) following *E. coli* stimulation shown as CD107a^+^, CD107a^+^GrzA^lo^, CD107a^+^GrzB^+^, and CD107a^+^Prf^+^. **(F, G)** MAIT cell pro-inflammatory cytokine production (IFNγ, TNF, and IL-17A) in healthy donors (n=5) and HAI patients (n=7). Box and whisker plots show median and the interquartile range. Statistical significance was calculated using Mann-Whitney U test. **** p<0.0001, *** p<0.001,** p<0.01,* p<0.05. ns, not significant. Gnly, granulysin; GrzA, granzyme A; GrzB, granzyme B; HAI, patients with healthcare-associated infections; HC, healthy control; IFN, interferon; IL, interleukin; Prf, perforin; TNF, tumor necrosis factor.

### Combined effects of TCR and cytokine signals on MAIT cell proliferation and cytolytic protein expression

Given the impaired MAIT cell functionality observed in HAI patients in this study, defining conditions that restore proliferative and cytotoxic potential is of clinical interest. Peripheral blood MAIT cells from healthy individuals were initially expanded *in vitro* using the MR1 ligand 5-OP-RU in combination with individual cytokines at their optimal concentrations, including IL-1β, IL-12, IL-15, IL-18, and IL-23 (S2A – S2D Figures). These cytokines are often elevated during infection and have the capacity to activate MAIT cells [10,20–25]. After 12 days, significant MAIT cell expansion was observed across all cytokine conditions (Figures 2A and 2B, S2C – S2E Figures) accompanied by concomitant upregulation of key cytolytic protein expression (Figures 2C – 2E). Optimal cytolytic protein expression occurred when both the MR1 ligand and cytokines were concurrently administered (S2F-S2L Figures). Notably, IL-15 most effectively promoted proliferation and strong cytolytic protein expression, with particularly high Gnly levels (Figure 2B – 2E, S2E Figure). IL-15 also appeared to exert a similar effect in promoting the highest proportion of polyfunctional cells (Figure 2F). Within the total T cell pool, most of the cytolytic protein expression following such stimulations appeared to be within the MAIT cell population (S3A Figure).

**Figure 2.**
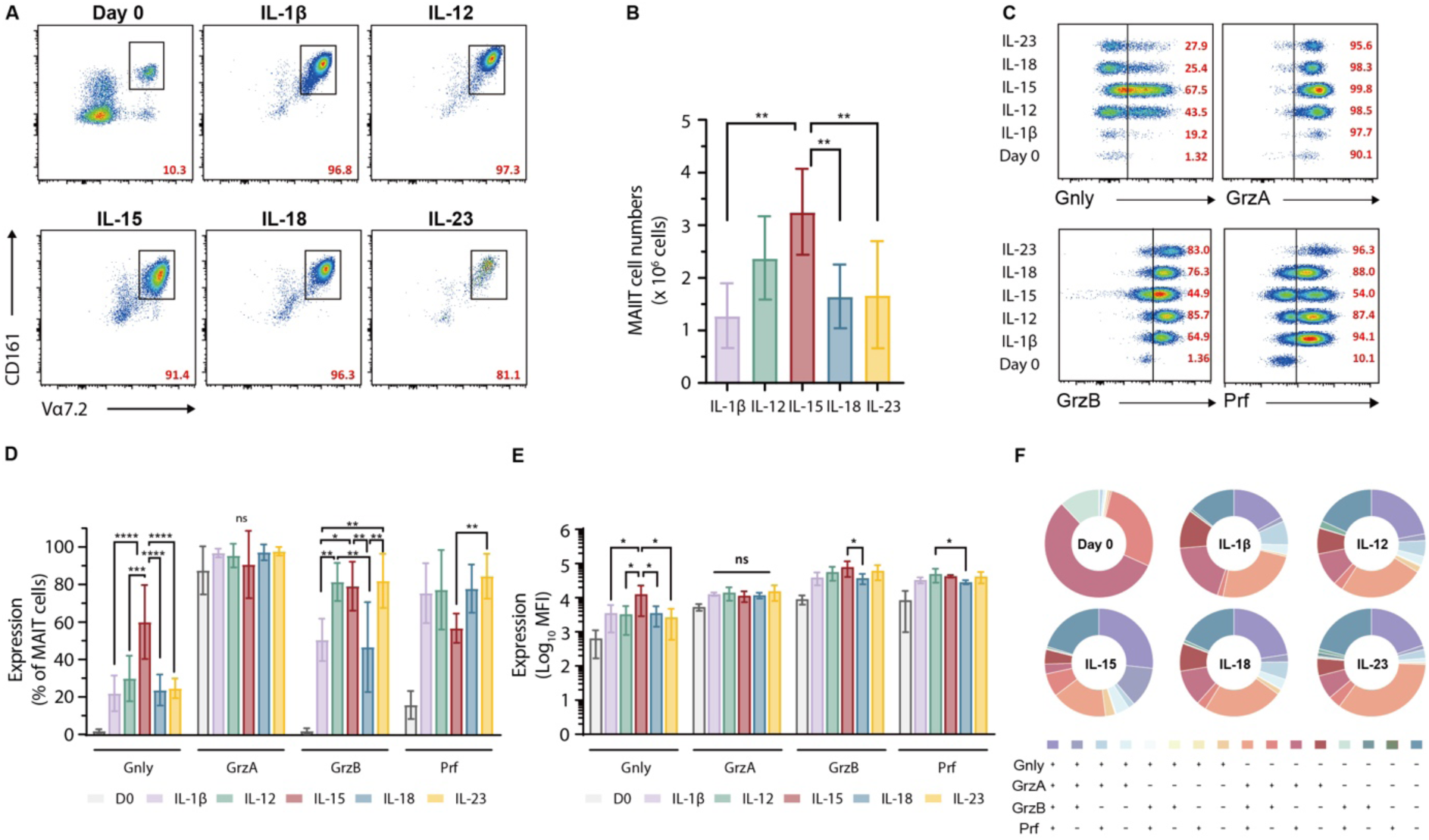
Modulation of MAIT cell proliferation and effector cytolytic protein expression in response to TCR and individual cytokine stimulations. **(A, B)** MAIT cell levels and **(C, D, E)** expression of Gnly, GrzA, GrzB, and Prf by MAIT cells following stimulation with the MR1 ligand 5-OP-RU with various single cytokines for 12 days (n=6). **(F)** Frequency of MAIT cell population expressing the indicated cytolytic proteins (n=6-8). Bar graphs show the mean and standard deviation. Data in the pie chart are presented as the mean. Statistical significance across multiple groups was calculated using one-way ANOVA followed by Tukey’s multiple comparison test. **** p<0.0001, *** p<0.001,** p<0.01,* p<0.05. ns, not significant. Gnly, granulysin; GrzA, granzyme A; GrzB, granzyme B; IL, interleukin; Prf, perforin.

We next evaluated how distinct cytokine combinations modulated MAIT cell cytotoxic profiles. Combinations tested included IL-1β+IL-23, IL-12+IL-18, IL-2+IL-7, and IL-2+IL-7+IL-15, based on prior evidence of such cytokine combinations in enhancing MAIT cell functionality [34–37]. All combinations supported MAIT cell proliferation and strong expression of multiple cytolytic proteins, with IL-2+IL-7 inducing the greatest MAIT cell expansion (Figure 3A – 3E, S3B Figure). All cytokine combinations appeared to induce MAIT cell cytolytic protein polyfunctional expression to similar extents (Figure 3F). Addition of IL-15 to IL-2+IL-7 did not appear to provide a further boost to MAIT cell proliferation and cytolytic expression (Figure 3A – 3E), although IL-15 modestly increased GrzB levels (Figure 3E). Altogether, these results show that both TCR and cytokine signals are required for effective MAIT cell proliferation and cytolytic protein expression, with IL-15 providing enhanced effects when used with MR1 ligand in the absence of other cytokines.

**Figure 3.**
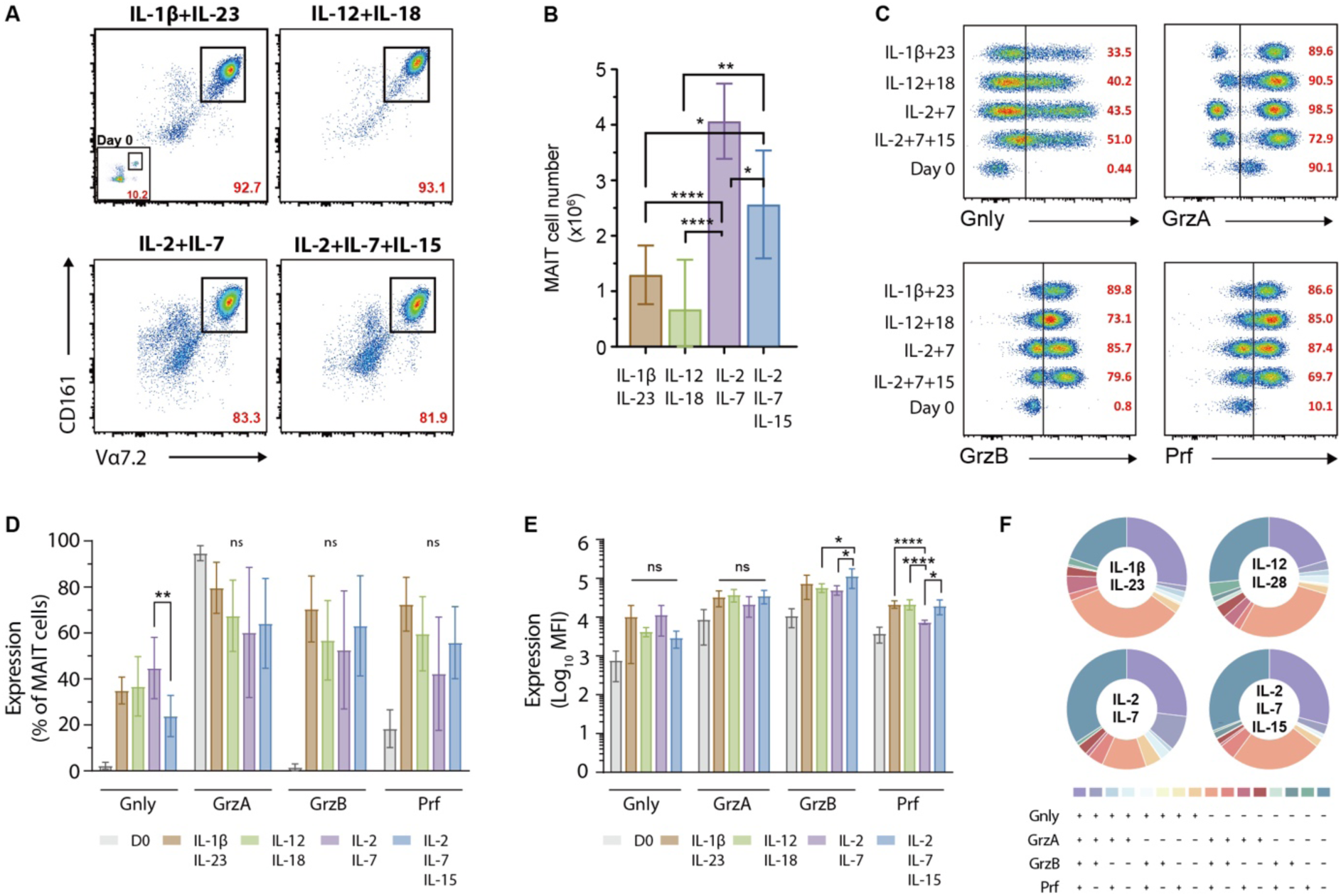
Modulation of MAIT cell proliferation and effector cytolytic protein expression in response to TCR and individual cytokine stimulations. **(A, B)** MAIT cell levels and **(C, D, E)** expression of Gnly, GrzA, GrzB, and Prf by MAIT cells following stimulation with the MR1 ligand 5-OP-RU with combined cytokines for 12 days (n=5-7). **(F)** Frequency of MAIT cell population expressing the indicated cytolytic proteins (n=5-7). Bar graphs show the mean and standard deviation. Data in the pie chart are presented as the mean. Statistical significance across multiple groups was calculated using one-way ANOVA followed by Tukey’s multiple comparison test. **** p<0.0001, *** p<0.001,** p<0.01,* p<0.05. ns, not significant. Gnly, granulysin; GrzA, granzyme A; GrzB, granzyme B; IL, interleukin; Prf, perforin.

### IL-15-expanded MAIT cells exhibit potent MR1-dependent cytotoxicity

Building on the observed proliferative and cytolytic responses induced by TCR and single cytokine co-stimulation, MAIT cell cytotoxic function were assessed using co-culture assays with target cells pulsed with the MR1 ligand 5-OP-RU (S3C Figure). All TCR and single cytokine co-stimulated MAIT cells demonstrated degranulation, as indicated by upregulation of the cytolytic exocytosis marker CD107a/LAMP-1 (Figure 4A). This degranulation was accompanied by apoptosis of the MR1 ligand-pulsed target cells (Figure 4B). Consistent with its ability to drive high cytolytic protein expression, IL-15-expanded MAIT cells exhibited stronger degranulation and more efficient killing of 5-OP-RU-pulsed targets (Figure 4A and 4B). Blocking MR1 significantly reduced both MAIT cell degranulation and target cell death, confirming MR1-dependent cytotoxicity (Figures 4A and 4B).

**Figure 4.**
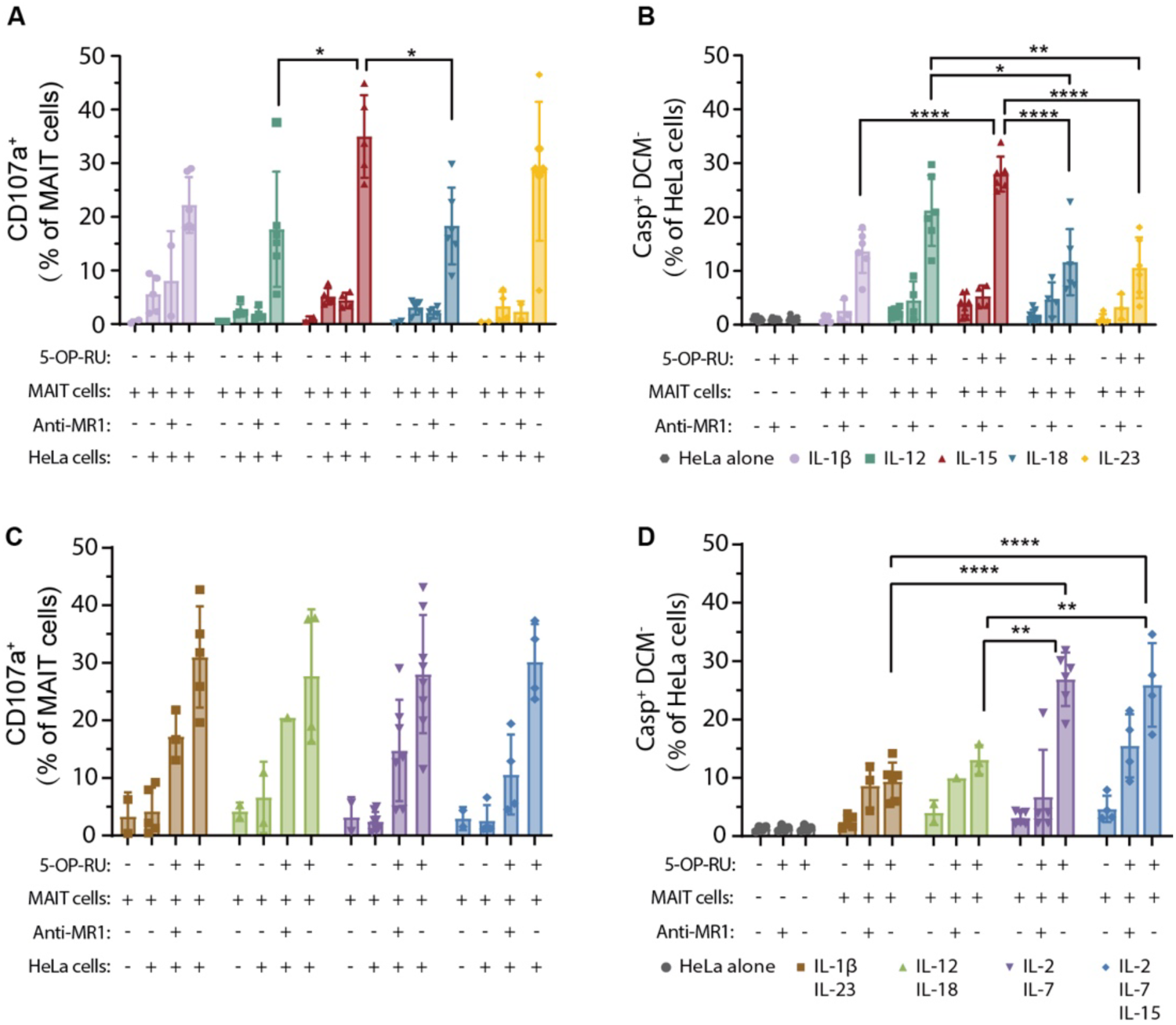
Effect on MAIT cell effector cytolytic function in response to TCR and various cytokine stimulations. **(A-D)** MAIT cell degranulation **(A, C)** and target cell killing, as indicated by active Caspase 3 staining **(B, D)**, following co-culture with 5-OP-RU–pulsed HeLa cells in the presence of anti-MR1 mAb or isotype control. MAIT cells were pre-expanded under either single **(A, B)** or combined cytokine **(C, D)** stimulation conditions. Bar graphs show the mean and standard deviation. Statistical significance across multiple groups was calculated using one-way ANOVA followed by Tukey’s multiple comparison test. **** p<0.0001, *** p<0.001,** p<0.01,* p<0.05. ns, not significant. 5-OP-RU, 5-(2-oxopropylideneamino)-6-D-ribitylaminouracil; DCM, amine-reactive live/dead cell marker; IL, interleukin; MR1, MHC-I related protein.

When evaluating cytokine combinations, MAIT cells expanded under tested conditions promoted MR1-dependent degranulation and cytotoxic function, with minimal differences observed for degranulation between cytokine groups (Figures 4C and 4D). Interestingly, IL-2 and IL-7 combination promoted the strongest killing of 5-OP-RU-pulsed target cells, independent of IL-15 addition (Figure 4D). The overall cytotoxic activity correlated with the extent of MAIT cell degranulation irrespective of stimulation conditions (S3D and S3E Figures). In summary, these findings indicate that IL-15, in combination with TCR stimulation, is sufficient to promote robust MAIT cell cytotoxicity.

### Functional recovery of MAIT cells from HAI patients following antigen and cytokine stimulation

Having identified conditions that enhance MAIT cell proliferation and cytotoxicity in healthy donors, their ability to restore MAIT cell function in HAI patients (Figure 1) was subsequently evaluated. Due to limited cell availability from patient samples, the IL-2+IL-7 combination was selected as a representative stimulation condition, based on the results showing strong proliferative and cytolytic responses (Figures 3A – 3E). Following expansion with 5-OP-RU and IL-2+IL-7, MAIT cells from HAI patients exhibited comparable proliferative capacity to those from healthy controls, as indicated by similar frequency and fold increases in cell numbers (Figures 5A – 5C). These expanded MAIT cells also upregulated key cytolytic effectors to levels approaching or matching those of healthy donors (Figures 5D – 5E). Cytotoxicity was evaluated through co-culture with 5-OP-RU-pulsed or *E. coli*-fed HeLa cells, where expanded MAIT cells from both healthy and HAI donors displayed potent degranulation and killing of target cells (Figures 5F – 5G). Together, these findings demonstrate that, despite numerical and functional impairments at baseline, MAIT cells from HAI patients can be functionally restored under defined stimulatory conditions. The recovery of cytotoxic function through IL-2+IL-7 and antigen stimulation supports the notion that MAIT cell function can be restored in the immunocompromised or HAI individuals.

**Figure 5.**
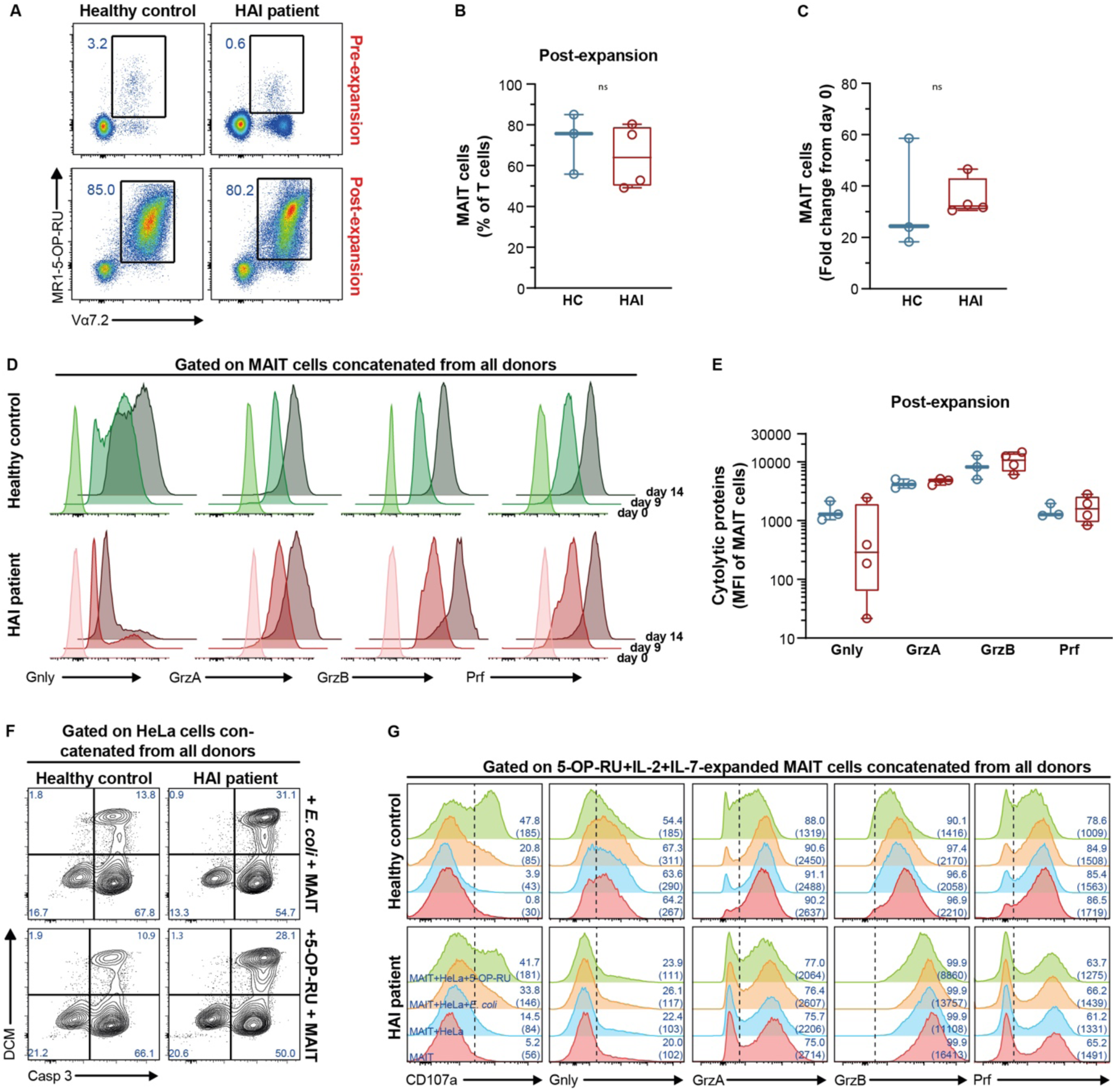
MAIT cell proliferation and functional restoration of cytolytic activity in residual cells from HAI patients following combined TCR and cytokine stimulation. **(A-E)** MAIT cell expansion **(A-C)** and expression of Gnly, GrzA, GrzB, and Prf **(D-E)** following 5-OP-RU stimulation in the presence of IL-2 and IL-7 for 17 days from healthy donors (n=3) and HAI patients (n=4). (F-G) Killing of the HeLa target cells (F) and MAIT cell degranulation and expression of cytolytic proteins following co-culture with 5-OP-RU–pulsed or *E. coli*-fed HeLa cells. MAIT cells from healthy donors (n=3) or HAI patients (n=4) were pre-expanded under 5-OP-RU and IL-2 + IL-7 stimulation conditions. Box and whisker plots show median and the interquartile range. 5-OP-RU, 5-(2-oxopropylideneamino)-6-D-ribitylaminouracil; Casp 3, active Caspase 3; DCM, amine-reactive live/dead cell marker; Gnly, granulysin; GrzA, granzyme A; GrzB, granzyme B; Prf, perforin.

### Secretomes from MR1 ligand and cytokine-stimulated MAIT cells exhibited potent antimicrobial efficacy against *E. coli* that produce carbapenemase

Our previous work demonstrated that MAIT cells respond to carbapenem-resistant *E. coli* (CREC) clinical isolates by secreting cytolytic proteins that enhance antimicrobial activity and restore carbapenem efficacy against extracellular CREC [18]. Here, we found that the antimicrobial response against *E. coli* persists even when these clinical CREC isolates [2,18] are cultured under carbapenem pressure (S4A Figure). This suggests that riboflavin-derived MR1 ligands are produced by CREC despite antibiotic exposure and that MAIT cells remain capable of detecting and responding to bacterial infection during antibiotic treatment. Having shown that MR1 ligand and cytokine co-stimulation differentially enhance MAIT cell cytolytic protein expression and cytotoxic function (Figurea 2A – 2E), we next aimed to identify cytokine conditions that effectively suppress bacterial growth and restore carbapenem efficacy. To this end, we engineered the *E. coli* BL21 strain to express the clinically relevant carbapenemases *bla_NDM-1_*, *bla_KPC-2_*, or *bla_OXA-48,_* representing class B, A, and D carbapenemases, respectively (S4B Figure). The activity of the transduced carbapenemases was assessed by monitoring imipenem hydrolysis, which generates acid and lowers the pH of the medium, as indicated by a color change in the phenol red indicator (S4C Figure). When cultured in the presence of various β-lactam antibiotics, including carbapenems, these engineered strains exhibited markedly elevated minimum inhibitory concentrations (MICs), indicating that the expression of the respective carbapenemases conferred functional resistance to carbapenems (S2 Table). MAIT cell secretomes collected from the different stimulation conditions were then tested for antimicrobial activity in the presence of imipenem against each of these engineered strains.

Cytolytic protein profiling of MAIT cell secretomes indicated that the production of multiple cytolytic proteins occurred at variable levels depending on the cytokine condition. Notably, MAIT cells stimulated with IL-15 produced significantly higher levels of GrzA, GrzB, and Prf compared to those stimulated with IL-1β or IL-18 (Figures 6A – 6D), although overall Prf secretion remained low (Figure 6D). Combinatorial cytokine stimulations did not significantly enhance cytolytic protein secretion beyond individual cytokine effects (S4D – S4G Figures).

**Figure 6.**
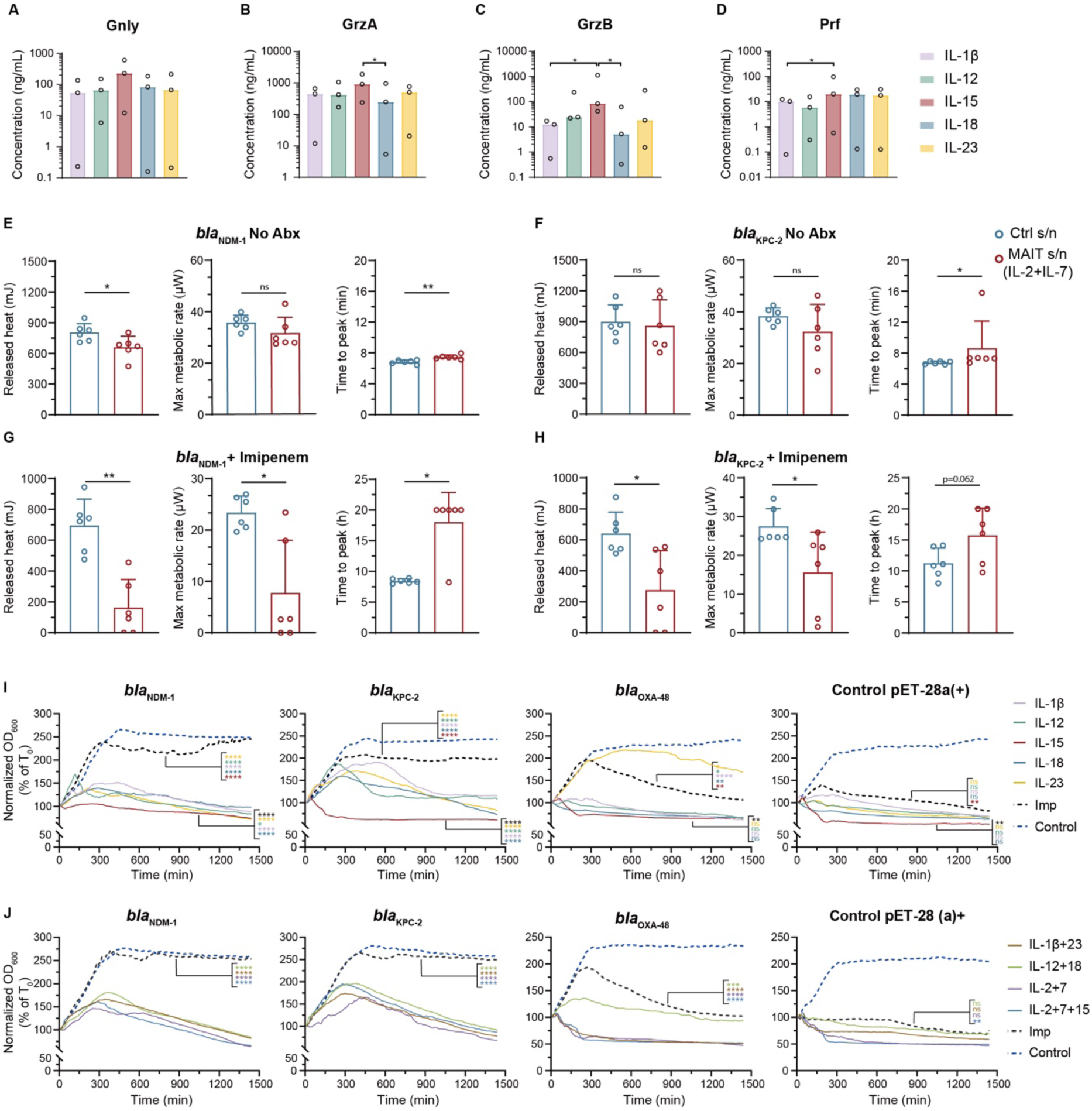
Effects of cytolytic protein–containing MAIT cell secretomes on the metabolism and growth of *E. coli* engineered to express carbapenemases. **(A-D)** Cytolytic protein levels within MAIT cell secretomes following stimulation with 5-OP-RU and single cytokines for 12 days (n=3). **(E-H)** Measurement of metabolic activity in *E. coli* expressing *bla*_NDM-1_ and *bla*_KPC-2_ carbapenemases following co-culture with MAIT cell secretomes, with or without imipenem, using isothermal microcalorimetry (n=5-6). **(I, J)** Growth of *E. coli* expressing *bla*_NDM-1_, *bla*_KPC-2_, and *bla*_OXA-48_ carbapenemases, along with control *E. coli*, in the presence of 50% minimum inhibitory concentration of imipenem for each strain, with or without MAIT cell secretomes derived from 12 days of expansion with 5-OP-RU and either single **(I)** (n=3) or combined **(J)** cytokines (pET-28 control n=2, all other conditions n=4). Bacterial growth was monitored by measuring the optical density at 600 nm (OD₆₀₀) over 24 hours and expressed as a percentage of the initial optical density at time zero. Bar graphs show the mean and standard deviation, whereas data presented as line graphs represent the mean. Statistical significance was calculated using the Mann-Whitney U test. Statistical significance across multiple groups was calculated using one-way ANOVA followrd by Tukey’s multiple compatison test **(A)**, or two-way ANOVA followed by Dunnett’s multiple comparison test **(I-J)**. **** p<0.0001, ** p<0.01,* p<0.05. ns, not significant. Abx, antibiotic; *bla*, β-lactamase; Gnly, granulysin; GrzA, granzyme A; GrzB, granzyme B; Imp, imipenem; KPC, *Klebsiella pneumoniae* carbapenemase; NDM, New Delhi metallo-β-lactamase; OXA, oxacillinase-group; Prf, perforin.

Isothermal microcalorimetry experiments were performed to confirm that MAIT cell secretomes enhance carbapenem activity. IL-2 and IL-7 were selected as the representative cytokine combination based on their ability to drive strong MAIT cell proliferation and cytotoxic polyfunctionality (Figures 3A – 3F), and our finding that IL-2+IL-7–stimulated MAIT cell secretomes restore carbapenem efficacy against extracellular carbapenem-resistant *E. coli* [18]. Secretomes from IL-2+IL-7–stimulated MAIT cells modestly reduced the metabolic output of *bla_NDM-1_* and *bla_KPC-2_* strains (Figures 6E – 6F). The effect was markedly enhanced when combined with imipenem, resulting in significantly reduced heat release, lower peak metabolic rate, and delayed time-to-peak kinetics (Figures 6G – 6H). These findings are consistent with our previous data, which showed that MAIT cell secretomes kill carbapenem-resistant *E. coli* by restoring carbapenem activity [18]. Additionally, these results indicate the capacity of MAIT cells to severely disrupt bacterial metabolism in the presence of carbapenem antibiotics.

To evaluate the direct antimicrobial effects, bacterial growth kinetics in the presence of imipenem, with and without MAIT cell secretomes, were monitored over 24 hours. MAIT cell secretomes from all cytokine conditions reduced bacterial growth in the presence of sub-inhibitory concentration of imipenem across all three resistant strains (Figure 6I, S5 Figure). Notably, secretomes derived from IL-15–stimulated MAIT cells exhibited the most rapid and sustained bacterial suppression (Figure 6I, S5 Figure), while combinatorial cytokines did not further enhance this effect (Figure 6J, S6 Figure). Moreover, time-kill studies confirmed that bacterial killing occurred rapidly, as IL-15-stimulated MAIT cell secretomes in combination with imipenem killed all tested *E. coli* strains expressing *bla_NDM-1_*, *bla_KPC-2_*, or *bla_OXA-48_* within 2 hours, with no regrowth detected over 24 hours (S7 Figure). Similar rapid killing was observed using secretomes from MAIT cells stimulated with cytokine combinations in the presence of imipenem (S8 Figure). In contrast, imipenem alone or control supernatants failed to clear the bacteria, indicating a combined effect of MAIT cell-derived cytolytic proteins and carbapenems. *E. coli* BL21 engineered to express *bla_OXA-48_* remained susceptible to imipenem in the absence of MAIT cell secretomes, consistent with the fact that OXA-48 is a relatively weak carbapenemase and does not typically confer high-level carbapenem resistance without additional resistance mechanisms.

## DISCUSSION

This study demonstrates that secreted factors from activated MAIT cells can restore the antimicrobial efficacy of imipenem against carbapenemase-producing *E. coli*. Activation with the cognate antigen together with γ-chain cytokines, particularly IL-15, led to robust MAIT cell expansion and cytolytic activity. The resulting secretomes restored imipenem efficacy against *E. coli* engineered to overexpress clinically relevant carbapenemases. The rapid bacterial clearance observed with MAIT cell secretomes in the presence of imipenem indicates a synergistic interaction. This synergy highlights the potential of γ-chain cytokines as effective MAIT cell activators in this setting. Notably, this strategy also enabled robust expansion and recovery of MAIT cells from patients with healthcare-associated infections (HAI), whose baseline populations were numerically and functionally impaired, resulting in strong proliferative capacity and regained cytolytic activity comparable to those of healthy donors. Collectively, these findings provide a translational framework for generating potent MAIT cells *ex vivo* from HAI and immunocompromised patients and support their application in host-directed strategies to enhance antibiotic efficacy and combat multidrug-resistant infections.

Building on our previous work demonstrating that MAIT cells suppress carbapenem-resistant CR-GNB through cytolytic mechanisms [18], the present study defines the key activation signals that most effectively potentiate these antimicrobial responses. We show that both MR1-dependent TCR engagement and cytokine stimulation are required for maximal MAIT cell cytolytic activity, as neither stimulus alone induced strong effector responses [19,20,38–40]. In combination, however, 5-OP-RU and γ-chain cytokines markedly promoted cytolytic programming of MAIT cells, with robust expression of Gnly, GrzA, GrzB, and Prf. Among cytokines tested, IL-15 most consistently supported proliferation and broad cytolytic protein co-expression, while the IL-2 plus IL-7 combination reproduced the strong proliferative and cytotoxic effects observed previously [18,37]. In contrast, IL-1β, IL-12, or IL-18 alone provided minimal additional benefit. Functional assays confirmed that culture with MR1 ligand and IL-15, or MR1 ligand and IL-2 plus IL-7, produced the highest MAIT cell killing activity. These findings support a strategy in which cognate antigen-TCR engagement together with IL-15, or IL-2 plus IL-7, optimally potentiates MAIT cell cytolytic function. Consistent with the activity in this setting, IL-15 is under evaluation in multiple clinical trials for cancer and chronic viral infections due to its ability to expand and activate cytolytic lymphocytes [41,42]. Optimal cytokine combinations and timing may vary by donor or clinical context and warrant further investigation.

MAIT cells in the current HAI patient cohort were both numerically depleted and functionally impaired. Residual MAIT cells exhibited markedly reduced expression of the cytolytic effectors Gnly and GrzB, key and interdependent mediators of bacterial killing [18,28], and produced less IFNγ and TNF upon stimulation. The combined reduction in frequency and function suggests a compromised ability to mount effective antimicrobial responses, including against AMR pathogens. Whether these deficits reflect pre-existing dysfunction associated with comorbidities [4] or redistribution of MAIT cells to infected tissues [10,43] remains unresolved, and follow-up studies after recovery may help address this important question. Nevertheless, our study importantly indicates that activation with the MR1 ligand 5-OP-RU plus γ-chain cytokines, including IL-2, IL-7, and IL-15, promoted robust MAIT cell proliferation and cytolytic potential. Applied to HAI patient samples, this strategy led to successful *ex vivo* MAIT cell expansion, restored cytolytic protein expression, and enabled effective killing of 5-OP-RU–pulsed targets and *E. coli*–infected cells, suggesting that even severely impaired MAIT cells remain amenable to functional rescue under optimal stimulation conditions.

MAIT cell secretomes generated under optimal activation suppressed *E. coli* expressing individual carbapenemases (*bla*_NDM-1_, *bla*_KPC-2_, or *bla*_OXA-48_) and restored imipenem efficacy against these resistant strains. IL-15 produced the strongest cytolytic protein secretion and yielded secretomes with the most rapid and sustained antimicrobial effects, whereas IL-2 plus IL-7, despite supporting proliferation, did not further enhance killing beyond IL-15. Importantly, time-kill assays showed that MAIT cell secretomes not only inhibited growth but directly killed carbapenemase-expressing *E. coli* in the presence of imipenem, confirming their bactericidal activity. Isothermal microcalorimetry further demonstrated marked reductions in bacterial metabolic activity, indicating further functional impairment beyond the membrane damage mediated by Gnly and GrzB [18]. The use of engineered *E. coli* strains allowed controlled assessment of secretome efficacy against defined resistance mechanisms, avoiding confounders present in clinical isolates with multiple resistance determinants, such as the co-expression of other β-lactamases, porin loss, and efflux pumps. Consistent with our prior work in CREC strains [18], these findings indicate that MAIT cell–derived Gnly and GrzB can bypass carbapenemase-mediated resistance, enabling carbapenems to regain potency. Overall, the findings from these studies suggest MAIT cell secretomes display broad antimicrobial activity across diverse resistance mechanisms.

In summary, we have defined a strategy to enhance MAIT cell antimicrobial function and contribute to restoring the efficacy of carbapenems against resistant bacteria. These results support the potential of MAIT cell-based therapy as an adjunctive treatment alongside conventional antibiotics in addressing antimicrobial resistance. From a translational perspective, the ability to expand functional MAIT cells from HAI patients using defined stimulation conditions raises the possibility to develop adoptive cell therapy approaches for AMR infections. In addition, this strategy could potentially be adapted to increase MAIT cell numbers and enhance function in at-risk individuals as a prophylactic measure. While further studies are needed to assess durability, scalability, and safety, these findings offer a possible direction for future host-directed interventions against multidrug-resistant bacterial infections.

## MATERIALS AND METHODS

### Ethics statement

Human peripheral blood samples were collected after obtaining informed written consent from all donors, in accordance with study protocols that conformed to the provisions of the Declaration of Helsinki. Ethical approvals were provided by the National University of Singapore Institutional Review Board (NUS-IRB reference codes B-15-088 and H-18-029), Singapore General Hospital Institutional Review Board (protocols 2015-3045 and 2019-2645), and Tsinghua Shenzhen International Graduate School Institutional Review Board (Refs. 202171 and 202295).

### MAIT cell expansion and culture

PBMCs were isolated by density-gradient centrifugation (Ficoll-Paque PLUS, Cytiva) and immediately used or cryopreserved in liquid nitrogen. MAIT cell expansion was performed as described [18] (Figure S2A). Cytokine-specific–supplemented medium (S3 Table) was refreshed every other day, and MAIT cell purity, cytolytic proteins, and counts were assessed on days 7, 12, and 17. Viable cells (>70% purity) were used for downstream assays. In selected experiments, MAIT cells were magnetically enriched and cultured for 48 h with specified cytokines before collection.

### MAIT cell functional assay

MAIT cell secretome antimicrobial activity, activation, and cytotoxicity assays were performed as previously described [10,15,18,45,46]. For antimicrobial assays, overnight *E. coli* subcultures were washed and resuspended in MAIT-cell or control supernatants (25% v/v) at 10⁵ CFU/mL and incubated at 37 °C for 24 h. In selected experiments, secretomes were supplemented with imipenem at half-MIC. Bacterial growth was monitored by OD₆₀₀ every 10 min (TECAN), and aliquots collected at indicated time points were plated on Mueller-Hinton agar to quantify CFU.

For MAIT cell activation, PBMCs were stimulated with formaldehyde-fixed *E. coli* for 24 h, followed by intracellular staining of cytolytic proteins and cytokines. Monensin was added during the final 6 h for cytokine detection. CD107a antibody (S4 Table) was included at culture initiation to measure degranulation.

For cytotoxicity assays, HeLa cells were seeded in 96-well plates and pulsed with 2 nM 5-OP-RU. MAIT effector cells were labeled with CellTrace Violet and added at an effector to target (E:T) ratio of 5:1. CD107a antibody, with or without anti-MR1, was included at culture start. After 3 h at 37 °C, apoptosis in HeLa cells was assessed by active Caspase-3 and viability staining (S4 Table).

### Statistical analysis

Data sets were first assessed for normality. Statistically significant differences between samples were determined using the unpaired *t* test or Mann-Whitney’s U test for unpaired samples, and the paired *t* test or Wilcoxon’s signed-rank test for paired samples. The one-way or two-way analysis of variance (ANOVA) followed by the appropriate post hoc test was used to detect differences across multiple samples. Correlations were assessed using the Spearman rank correlation. Two-sided *p* < 0.05 were considered significant. Statistical analyses were performed using Prism software version 10.0.2 (GraphPad).

## Acknowledgments

We thank all donors, healthcare personnel, study coordinators, administrators, and laboratory managers involved in this work. The MR1 tetramer technology was developed jointly by J. McCluskey, J. Rossjohn, and D. Fairlie, and the material was produced by the NIH Tetramer Core Facility as permitted to be distributed by the University of Melbourne.

## Funding

Science, Technology and Innovation Commission of Shenzhen Municipality grant WDZC20220819153248002 (EL)

Tsinghua Shenzhen International Graduate School grant QD2022018C

(EL) Tsinghua Shenzhen International Graduate School grant JC2022007 (EL)

Swedish Research Council Grant 2015-00174, Marie Skłodowska Curie Actions, Cofund, Project INCA 600398 (EL)

NMRC Collaborative centre grant NMRC/CG/C005B/2017_SGH (ALHK)

CoSTAR-HS ARG Seed Fund 2018/02 (ALHK)

National Natural Science Foundation of China grant W2433196 (AMK)

Science, Technology and Innovation Commission of Shenzhen Municipality grant WDZC20231130105009001 (AMK)

National Institutes of Health RO1 grant AI148407-01A1 (DF)

NHMRC Investigator grant 2009551 (DF)

Australian Research Council grant CE200100012 (DF)

## Author contributions

Conceptualization: ALHK, EL

Methodology: XY, AMK, WRS, FH, EL

Investigation: XY, AMK, WRS, FH, KLC, LH, ZL

Data analysis: XY, AMK, WRS, FH, NGC, EL

Provision of critical materials: JWYM, DPF, LFW, JKS, ALHK

Supervision: ALHK, EL

Writing – original draft: XY, AMK, WRS, EL

Writing – review and editing: DPF, JKS

All authors reviewed and approved the final manuscript.

## Competing interests

XY, FH, and EL are listed as inventors on a granted patent (No. ZL202211393019.8) filed with the China National Intellectual Property Administration and owned by Tsinghua Shenzhen International Graduate School. JYWM and DPF are named inventors on a granted patent (No. WO2015/149130, World Intellectual Property Organization) owned by University of Queensland, Monash University, and University of Melbourne. The rest of the authors declare that they have no competing interests associated with this study.

## Data and materials availability

All data required to evaluate the conclusions in the paper are present in the paper and/or the Supplementary Materials upon publication.

## Supplementary materials

Supplementary tables. Supplementary figures and legends.

Supplementary materials and methods.

